# Esterase Activity is Affected by Genetics, Age, Insecticide Exposure, and Viral Infection in the Honey Bee, *Apis mellifera*

**DOI:** 10.1101/415356

**Authors:** Frank D. Rinkevich, Joseph W. Margotta, Michael Simone-Finstrom, Lilia I. de Guzman, Kristen B. Healy

## Abstract

Non-target impacts of insecticide treatments are a major public and environmental concern, particularly in contemporary beekeeping. Therefore, it is important to understand the physiological mechanisms contributing to insecticide sensitivity in honey bees. In the present studies, we sought to evaluate the role of esterases as the source of variation in insecticide sensitivity. To address this question, the following objectives were completed: 1) Evaluated esterase activity among honey bee stocks, 2) Assessed the correlation of esterase activity with changes in insecticide sensitivity with honey bee age, 3) Established if esterases can be used as a biomarker of insecticide exposure, and 4) Examined the effects of Varroa mite infestation and viral infection on esterase activity.

Results indicated that honey bees have a dynamic esterase capacity that is influenced by genetic stock and age. However, there was no consistent connection of esterase activity with insecticide sensitivity across genetic stocks or with age, suggests other factors are more critical for determining insecticide sensitivity. The trend of increased esterase activity with age in honey bees suggests this physiological transition is consistent with enhanced metabolic rate with age. The esterase inhibition with naled but not phenothrin or clothianidin indicates that reduced esterase activity levels may only be reliable for sublethal doses of organophosphate insecticides. The observation that viral infection, but not Varroa mite infestation, reduced esterase activity shows viruses have extensive physiological impacts. Taken together, these data suggest that honey bee esterase activity toward these model substrates may not correlate well with insecticide sensitivity. Future studies include identification of esterase substrates and inhibitors that are better surrogates of insecticide detoxification in honey bees as well as investigation on the usefulness of esterase activity as a biomarker of pesticide exposure, and viral infection.

## Introduction

The history of honey bee kills upon contact with insecticides has been documented since the advent of modern insecticides [1]. Beekeeper surveys report that pesticide exposure significantly increases annual colony losses [2]. Considering that a number of insecticides used in agriculture and vector control exhibit high toxicity to honey bees and that honey bees regularly encounter numerous of pesticides within the colony [3], potential synergistic interactions among these pesticides [4] may contribute to poor colony health.

Insects possess an array of metabolic mechanisms such as esterases, cytochrome P450s, and glutathione-S-transferases to detoxify pesticides, plant allelochemicals, and other xenobiotics [5]. Esterases are a type of hydrolase that metabolizes compounds by cleaving the ester bonds of a substrate resulting in separate acid and alcohol products [6]. Quantitative increases [7] as well as qualitative changes in esterase activity [8] may lead to reduction in insecticide sensitivity. In honey bees, esterase expression and activity are upregulated in response to exposure to *p*-coumaric acid [9], coumaphos [10], thiamethoxam [11], deltamethrin, fipronil, and spinosad [12]. Esterase inhibitors significantly increase sensitivity to phenothrin [13], tau-fluvalinate, cyfluthrin [14], fenpyroximate, and thymol [15], suggesting that esterase-mediated detoxification significantly influences pesticide sensitivity. Therefore, understanding the factors that affect honey bee esterase activity may yield insight into differences in insecticide sensitivity.

A myriad of factors such as age, diet, and genetics may affect insecticide sensitivity [13, 16, 17], but little research has been done on the underlying physiological mechanisms. Therefore, we decided to investigate a number of common factors that previous work suggests may affect honey bee physiological processes with a particular focus on esterase activity.

The current study aimed to tease apart several factors that influence insecticide sensitivity and esterase activity in honey bees.

1) Esterase comparison among honey bee stocks. Earlier studies demonstrated that insecticide susceptibility varies among Italian, Russian, and Carniolan stocks of honey bees, and esterase inhibition has been shown to increase sensitivity to phenothrin [13]. This led us to hypothesize that esterases may contribute to variation in insecticide sensitivity among honey bee stocks and across age.

2) Changes in esterase activity with age. Because of changes in pesticide sensitivity occurring with increased age [13], we assessed esterase activity in worker bees of different ages to compare if changes in esterase activity correlated with changes in insecticide sensitivity.

3) Esterase inhibition by insecticides. Numerous sublethal effects of pesticides have been demonstrated [18-20], and esterase activity has been proposed as a biomarker of high levels of pesticide exposure [11, 12]. Therefore we assayed the changes of esterase activity upon exposure to experimentally-determined sublethal levels of the insecticides naled, phenothrin, and clothianidin.

4) Impacts of Varroa mite infestation and viruses on esterase activity. All honey bee colonies in the US are infested with the ectoparasitic mite, *Varroa destructor* (hereto referred to as the Varroa mite). Varroa mites and the associated viruses they transmit are among the most significant factors relating to colony failure [21, 22]. These factors were both tested because mite infestation and viral infection alone and in combination have multifactorial effects on honey bee physiology and response to insecticide activity [20, 23, 24]. Therefore, the effects of Varroa mite infestation and viral infection on esterase activity were also investigated. Results are discussed in terms of how progression through the honey bee’s life history and the impact of biotic factors influence esterase capacity. We further suggest the notion of developing esterases as a potential biomarker of insecticide exposure and viral infection.

## Materials and Methods

### Esterase comparison among honey bee stocks

#### Esterase Assays

The model substrates fo r esterase activity (1-naphthyl acetate (1NA), *para*-nitrophenyl acetate (PNPA)), Fast Blue B, sodium dodecylsulfate, and Bradford Reagent were obtained from Sigma (St. Louis, MO). 1NA and PNPA were used because they are model substrates that are representative of general esterase and cholinesterase activity, respectively [25].

Esterase activity was performed according to established protocols modified for a 96-well plate [26]. Bee abdomens were homogenized with a disposable pestle in 1 ml of 100 mM sodium phosphate buffer (pH 7.4) in a microcentrifuge tube. Samples were spun for 10 m at 4^°^C at 10,000g. The supernatant was diluted 1:10 in 100 mM sodium phosphate buffer (pH 7.4) for use in esterase and Bradford assays.

For 1NA endpoint assays, 20 ul of homogenate was added to a well of a 96-well plate (model 9017, Corning Life Sciences, Corning NY) in duplicate. Each well received 200 ul of 0.3 mM 1NA (final concentration, dissolved in 100 mM sodium phosphate buffer (pH 7.4)). Plates were held at room temperature (RT) for 15 min. Fifty ul of staining solution (0.15 g Fast Blue B dissolved in 15 ml distilled water and 35 ml of 5% (w/v) sodium dodecylsulfate), and color was allowed to develop for 5 m at room temperature. Plates were read at 570 nm in a Spectramax 190 with SoftMax Pro 7.0 software (Molecular Devices, Sunnyvale, CA). Standard curves were run in parallel each day with 2-fold serial dilutions of 1-naphthol.

The PNPA kinetic assay [27] was performed with 20 ul of enzyme homogenate added to a 96 well plate in duplicate. Control wells received 20 ul of 100 mM sodium phosphate buffer (pH 7.4). Just prior to the PNPA assay, 0.1 ml of 100 mM PNPA (dissolved in acetonitrile) was added to 9.9 ml of 50 mM sodium phosphate buffer (pH 7.4) and vortexed 5 s. Each well received 200 ul of the diluted PNPA solution (1 mM PNPA final concentration), and the changes in absorbance were immediately read every 10 s for 2 m in a Spectramax 190 at 405 nm. PNPA activity was calculated by subtracting the average control activity from the experimental samples.

Protein concentration was determined by the Bradford method [28] by placing 10 ul of supernatant into a 96 well plate in duplicate. Each well received 200 ul of diluted Bradford Reagent (BioRad, Hercules CA), incubated at room temperature for 5 m, and then absorbance was read in a Spectramax 190 at 595 nm. A standard curve was generated using serial 2-fold dilutions of bovine serum albumin. Esterase activity towards 1NA and PNPA was standardized by protein content.

### Honey bee colonies and collections

Italian, Carniolan, and Russian queens were purchased from commercial breeders. Colonies were established at the USDA-ARS Honey Bee Breeding, Genetics, and Physiology Lab in Baton Rouge, LA. All colonies were maintained using standard management practices with no miticide applications, antibiotic treatments, or supplemental feeding.

Frames of emerging adult worker bees were removed from colonies and held at 33±1^°^C, 70±5% RH in continuous darkness overnight. Newly emerged adult bees were sorted into groups of 20 in 475 ml wax paper cups and supplied with cotton balls soaked with 50% sucrose solution (w/v). These bees were held at the environmental conditions listed above until 3-days of age and then frozen at - 80^°^C until further use in esterase assays described above. A total of 30 bees (5 individuals from 6 colonies) for each of the 3 honey bee stocks were used in esterase assays. Esterase activity levels between stocks were compared with Wilcoxon-Rank Sum test with post-hoc multiple comparisons test (α=0.05) using JMP (SAS, Cary, NC).

### Changes in esterase activity with age

#### Colonies with normal demographics

Brood frames were removed from 6 colonies of Italian honey bees each consisting of 2 deep boxes with brood frames and 1 medium honey super with >30,000 worker bees. Adults were allowed to emerge overnight from brood frames, marked with a dot of enamel paint on the notum, and returned to their respective colonies. Marked bees were collected either from inside the hive or returning from a flight every 3 to 5 days up to 31-days of age. At least 5 bees were collected each sampling date from each colony. Samples were frozen at −80^°^C until used in esterase assays described above. The correlation of age and esterase activity was compared with Spearman’s Rank Correlation using JMP.

### Correlation of esterase activity with changes in insecticide sensitivity with age

Newly emerged adult bees were marked with enamel paint and returned to source colonies as described above. A total of 10 source colonies were used. Bees were collected at 3-, 14-, 21-, and 28-days of age in groups of 10 into wax paper cups covered with nylon tulle secured with a rubber band. Topical bioassays with phenothrin (98.4% purity, ChemService, West Chester PA) and naled (99.0% purity) were performed as previously described [13]. Bees were anaesthetized with CO_2_ for <30s and a 1 ul drop of insecticide was applied to the notum with a Hamilton repeating syringe. Control bees were treated with acetone. Treated bees were provided a cotton ball soaked with 50% sucrose solution and held in an incubator under the environmental conditions listed above. At least 1 rep at each dose was used from each colony on each collection day. A subsample of 8 bees was collected at each collection date and stored at −80^°^C until used in esterase assays described above. The LC_50_ was calculated using Probit analysis with Abbott’s correction for control mortality [29] in Minitab (State College, PA) and expressed in units of ug insecticide/bee. The correlation of esterase activity with age and insecticide sensitivity was compared with Spearman’s Rank Correlation using JMP.

### Insecticide inhibition of esterase activity

#### Clothianidin bioassay

Clothianidin is a neonicotinoid insecticide that is widely used as a seed treatment for corn and soy beans. It is frequently found in honey bee colonies and may have detrimental impacts on honey bees [30, 31]. The LC_50_ for clothianidin (99.5% purity, ChemService, West Chester PA) was determined by a feeding bioassay according to previously published methods [13]. Newly emerged adult Italian honey bees from 3 colonies were sorted into groups of 20 in 475 ml wax paper cups and held at 33±1^°^C with 70±5% RH in continuous darkness until 3-days of age. Bees were fed 50% sucrose solution (w/v) containing concentrations of clothianidin that produced >1% and <99% mortality *ad libitum* through a perforated microcentrifuge tube. Mortality was recorded after 24 h. The LC_50_ was calculated using Probit analysis with Abbott’s correction for control mortality [29] in Minitab (State College, PA) and expressed in units of ng clothianidin/ml.

### *In vivo* esterase inhibition

Newly emerged adult Italian honey bees from 6 colonies were collected and aged to 3-days of age as described above. An experimentally-determined maximum-sublethal treatment of phenothrin (0.1 ug/bee, topical [13]), naled (0.066 ug/bee, topical [13]), or clothianidin (2.15 ng/ml, feeding, from above experiment) was administered to these bees. To determine the dose-dependence of esterase inhibition, sublethal doses of naled (i.e., 0.05, 0.033, and 0.025 ug/bee) were applied to 3-day old bees in a separate experiment. Control bees were treated with 1 ul of acetone for control topical bioassays or 50% sucrose solution with 0.001% acetone for feeding bioassays. Bees were collected at 24 hours after treatment and frozen at −80^°^C until use in esterase assays described above. Esterase activity data from insecticide exposed bees were compared with Wilcoxon-Rank Sum Test with post-hoc multiple comparisons test (α=0.05) using JMP. The correlation of naled doses and esterase activity was compared with Spearman’s Rank Correlation using JMP.

### Impacts of Varroa mite and virus on esterase activity

#### Varroa mite infestation

Frames of emerging adults were removed from 4 colonies of Italian bees that showed no symptoms of viral infection. Emerging adults and the associated brood cells from which they emerged were examined for Varroa mite infestation by a single foundress [32]. Varroa mite infested and uninfested adults were collected into separate wax paper cups provisioned with a cotton ball saturated with 50% sucrose solution. These bees were held in an incubator at 33±1^°^C with 70±5% RH in continuous darkness until 3-days of age, then stored at −80^°^C until used in esterase assays as described above. Esterase activity from Varroa mite infested bees was compared with Wilcoxon-Rank Sum Test using JMP.

### Virus injection

Solutions of *Deformed wing virus* (DWV), *Chronic bee paralysis virus* (CBPV), and *Black queen cell virus* (BQCV) were semi-purified by grinding 10 symptomatic adult bees (DWV, CBPV) or larvae (BQCV) in phosphate buffered saline (PBS) and centrifuged in a 15 mL tube at 4,700g for 20 m at 4°C. The supernatant was then filtered through a 0.2 um syringe filter and maintained at until 4°C use within a week following standard protocols [33]. Viral titers were determined using standard curves generated from plasmid standards containing the sequence listed above (generated by GeneArt, Invitrogen). Linearized plasmid standards containing 10^5^ to 10^12^ copies per reaction were used as templates to assess primer efficiency and quantify the amount of virus following standard practices [34-36]. Linear standard equations were generated using the log_10_ of the initial plasmid copy number. The genomic region encoding a capsid protein for each of the viruses was as follows: DWV Forward— GAG ATT GAA GCG CAT GAA CA and Reverse— TGA ATT CAG TGT CGC CCA TA (AY292384.1, [37]); CBPV Forward— CGC AAG TAC GCC TTG ATA AAG AAC and Reverse—ACT ACT AGA AAC TCG TCG CTT CG (EU122229.1, [38]); BQCV Forward—TTT AGA GCG AAT TCG GAA ACA and Reverse— GGC GTA CCG ATA AAG ATG GA (HQ655494.1, [37]).

Newly emerged adult bees (<1 h) from 3 colonies were collected into a 20 ml scintillation vial and chilled on ice. Bees were injected between the dorsal abdominal tergites with 3 ul of semi-purified virus (DWV @ 10^7^ copies/ul; CBPV @ 10^4^ copies/ul; BQCV @10^7^ copies/ul) using a Hamilton syringe fitted with a 30G needle at an infusion rate of 1ul/sec using a Micro4TM Microsyringe Pump Controller adapted from standard methods [33, 39]. Two sets of control bees were either uninjected or injected with PBS. Adults were collected in to a wax paper cup provisioned with a cotton ball with 50% sucrose solution and held in an incubator at 33+1^°^C with 70+5% RH in continuous darkness until 3-days of age. Survivors were collected and stored at −80^°^C until used in esterase assays as described above. Esterase activity in virus-injected bees was compared via One-Way ANOVA with Tukey’s HSD post-hoc test using JMP.

## Results

### Esterase activity varies among honey bee stocks

Carniolan and Italian honey bees exhibited significantly higher esterase activity towards 1NA (x^2^=10.4, df=2, p<0.01, Fig 1) compared to Russian honey bees. Italian honey bees showed significantly higher esterase activity towards PNPA compared to both Carniolan and Russian honey bees, and Carniolan honey bees had significantly higher activity compared to Russian honey bees (x^2^=17.5, df=2, p<0.01, Fig 1).

**Fig 1.**
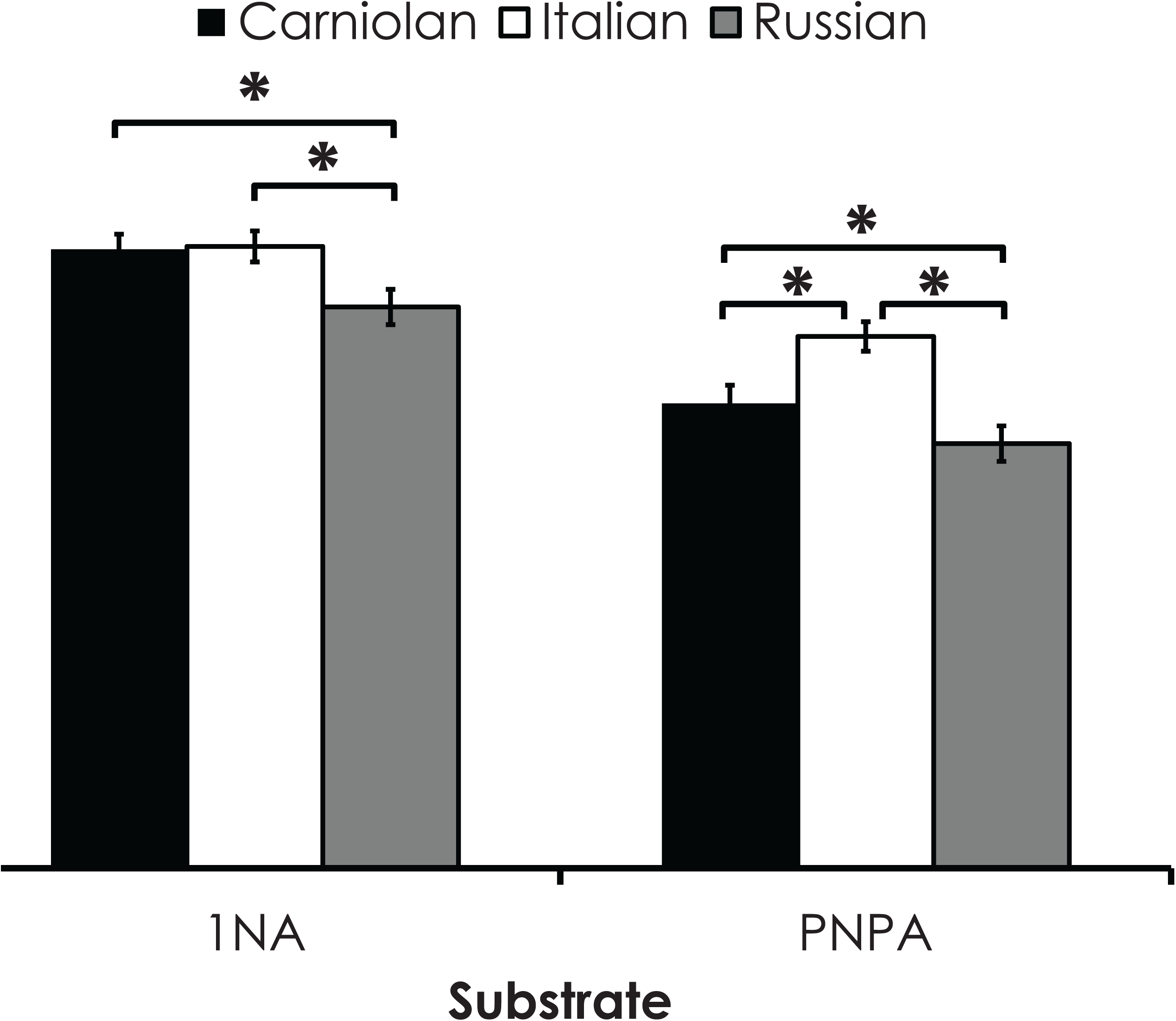
Esterase activity towards 1NA and PNPA from Carniolan, Italian, and Russian honey bees. The bar with an * indicate significant differences between stocks. Data are the average ± SEM.

### Changes in esterase activity with age

#### Esterase activity positively correlates with honey bee age

Esterase activity towards both 1NA and PNPA was significantly correlated with age in honey bees in colonies with normal demographics (1NA ρ=0.86, df=13 p<0.01, Fig 2A; PNPA ρ=0.91, df=13, p<0.01, Fig 2B).

**Fig 2.**
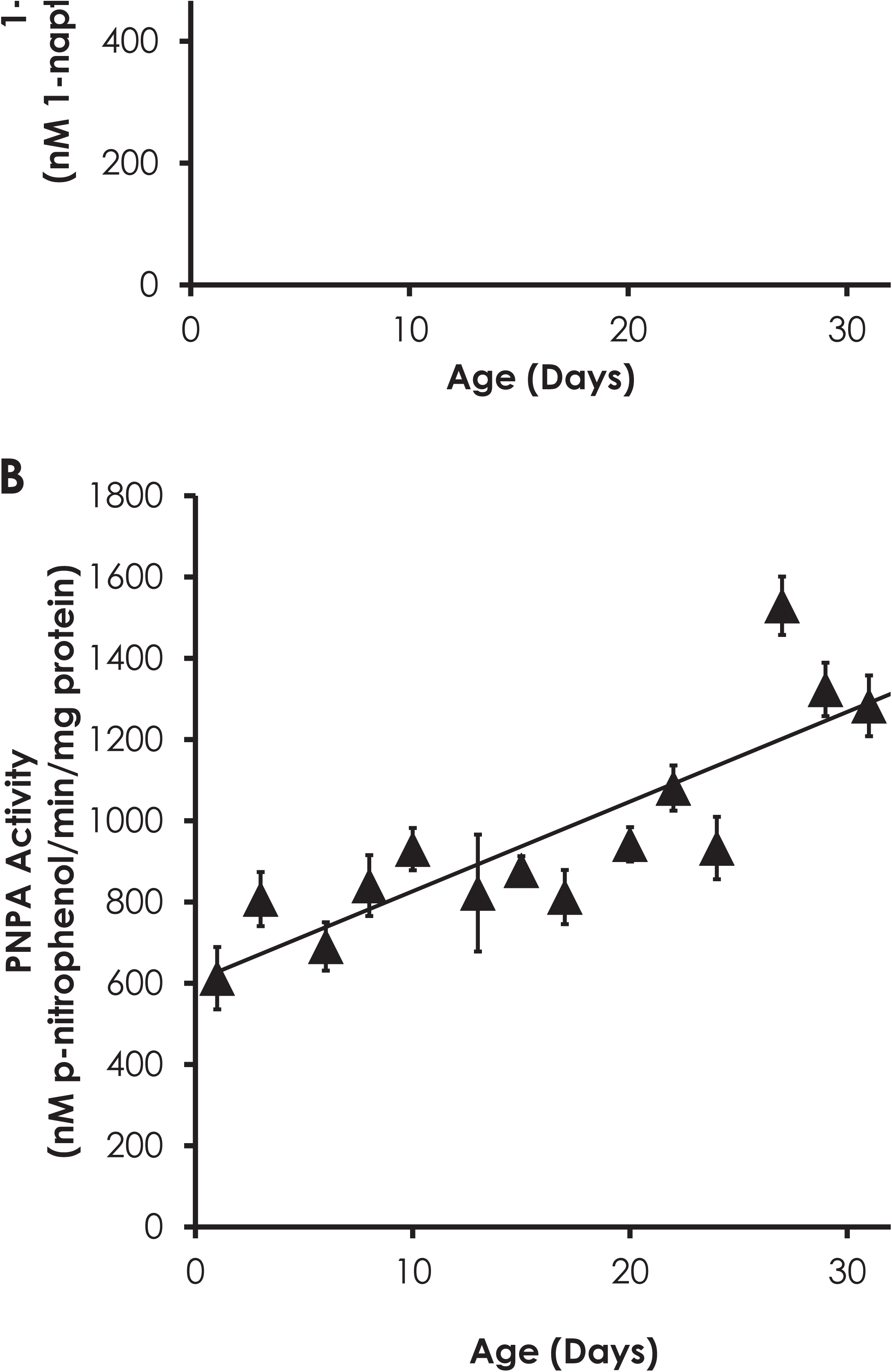
Changes in esterase activity with age. Esterase activity towards 1NA (A) and PNPA (B) significantly increased with age. Data are the average ± SEM.

### Changes in insecticide sensitivity with honey bee age do not correlate with esterase activity

Sensitivity to phenothrin did not significantly decrease with age in honey bees from colonies with normal demographics (ρ=0.80, df=3, p=0.20; Fig 3A), but older bees (i.e. 21- and 28-day old bees) were significantly less sensitive to phenothrin than younger bees (i.e. 3- and 14-day old bees). Naled sensitivity significantly increased with age in honey bees from colonies with normal demographics (ρ=-1.00, df=3, p<0.01; Fig 3B). Phenothrin sensitivity did not correlate with esterase activity towards 1NA (ρ=0.00, df=3, p=1.0; Fig 4A) or PNPA (ρ=0.80, df=3, p=0.20; Fig 4C). There was a significant negative correlation of naled sensitivity with esterase activity towards PNPA (ρ=-1.00, df=3, p=<0.01 Fig 4D), but not 1NA (ρ=-0.40, df=3, p=0.60 Fig 4B)

**Fig 3.**
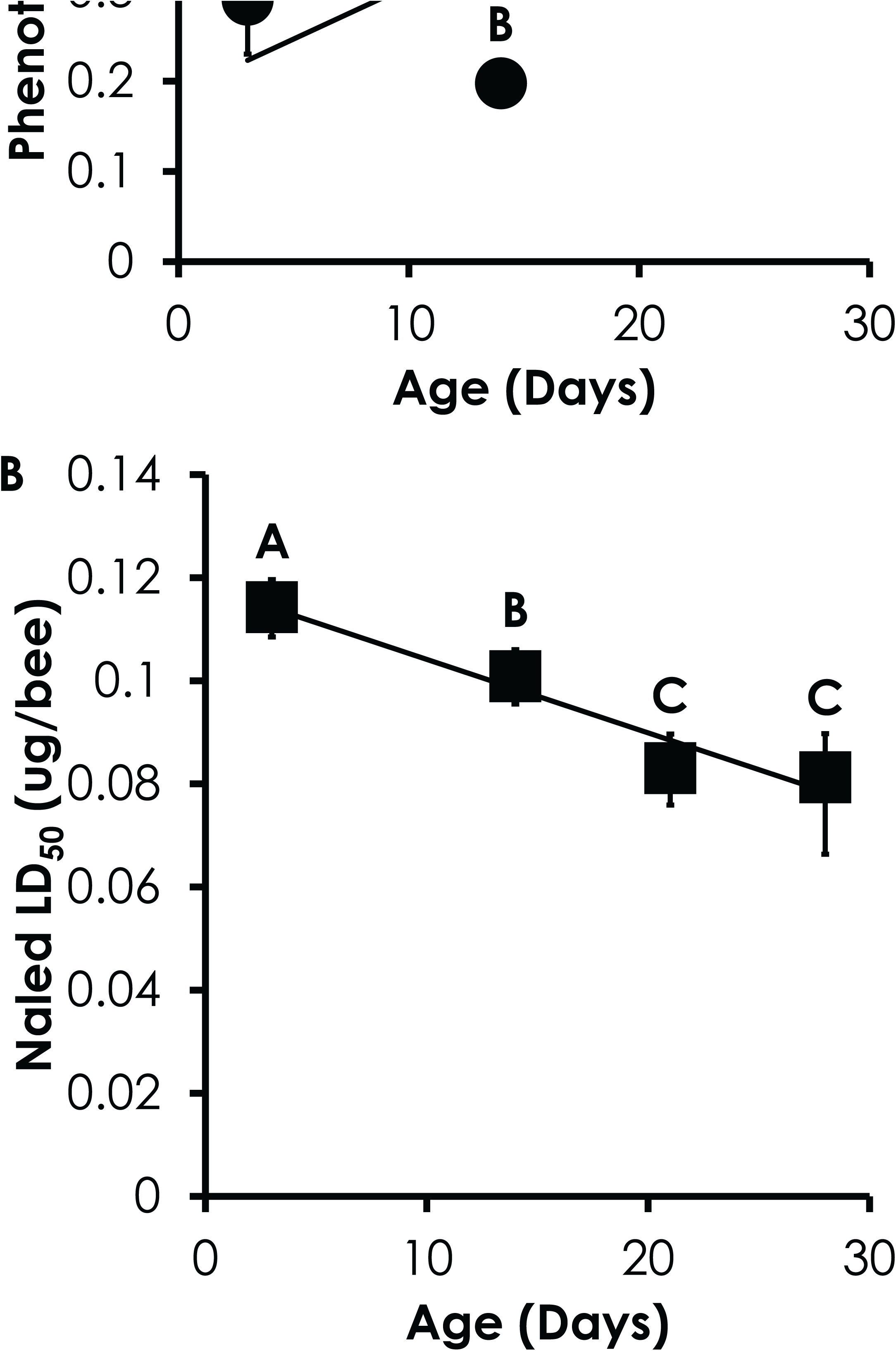
Changes in insecticide sensitivity with age in honey bees from colonies with normal demographics. Sensitivity to phenothrin decreased with age (A), while sensitivity to naled increased with age (B). Letters indicate significant differences in insecticide sensitivity at different ages. Data are the average ± 95% CI.

**Fig 4.**
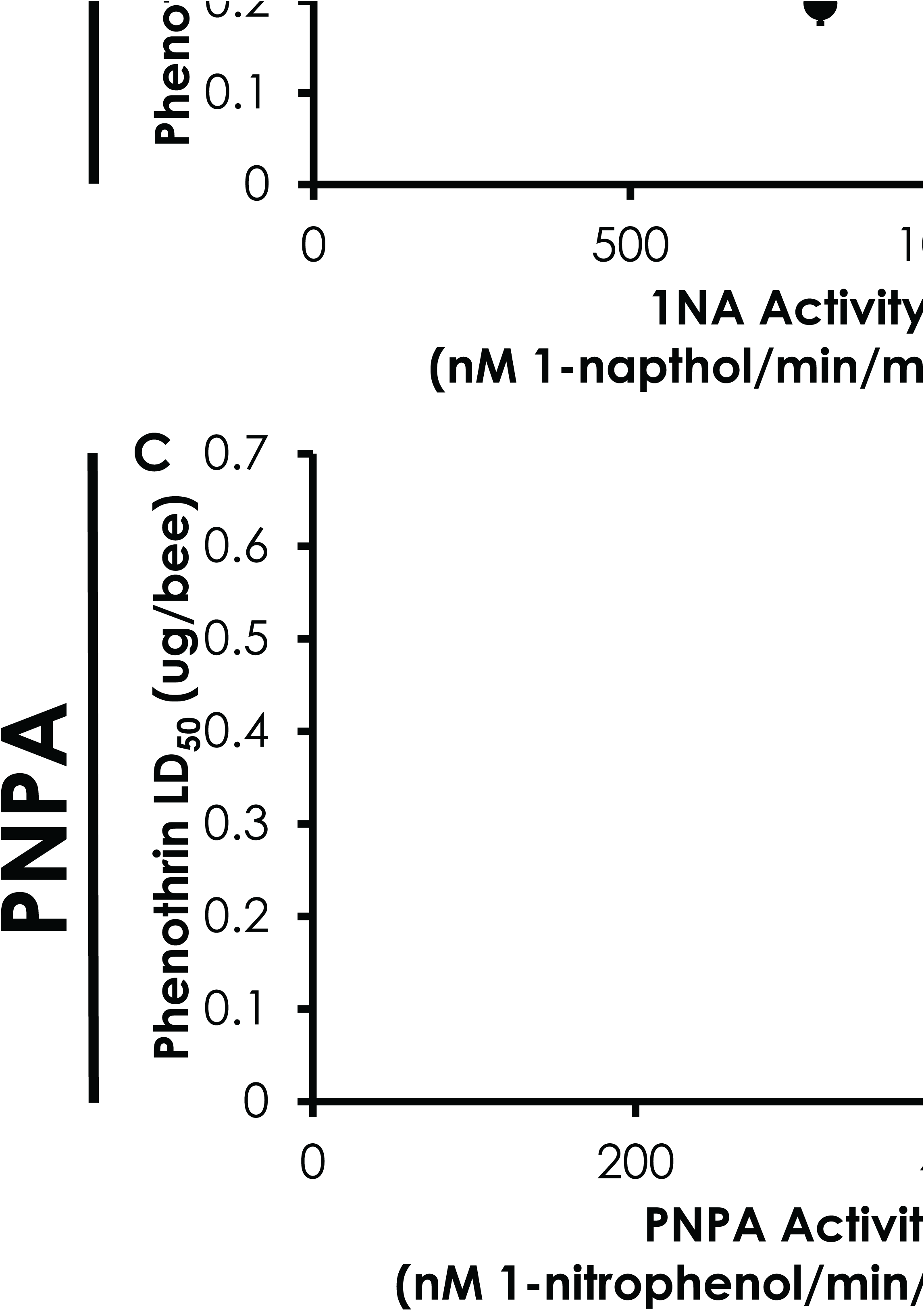
Correlation of esterase activity and insecticide sensitivity. Esterase activity towards 1NA was not correlated with sensitivity to phenothrin (A) or naled (B). There was no correlation of esterase activity towards PNPA and phenothrin sensitivity (C), but it was significantly correlated with naled sensitivity (D). Data are shown as the LD_50_ ± 95%CI.

### *In vivo* esterase inhibition by insecticides

#### Determination of maximum sublethal clothianidin concentration

The LC_50_ for clothianidin was 132.6 ng/ml (Table 1). The ratio of the LC_90_ to LC_10_ was 92-fold. Based upon these results, the maximum sublethal clothianidin concentration was calculated to be 2.1 ng/ml and verified by bioassays.

**Table 1.** Clothianidin toxicity to Italian honey bees. The LC values are expressed in ng clothianidin/ml sucrose solution. Values in parenthesis represent the 95% confidence interval and standard error for the LC values and slope, respectively.

| Compound | n | LC<sub>10</sub> | LC<sub>50</sub> | LC<sub>90</sub> | Slope |
| --- | --- | --- | --- | --- | --- |
| Clothianidin | 1052 | 13.8 (6.9-22.5) | 132.6 (103.9-160.3) | 1276.2 (986.4-1813.2) | 1.3+(0.1) |

### Sublethal insecticide exposure in vivo esterase inhibition varies with insecticide class

Both 1NA and PNPA activities were significantly inhibited by application of sublethal dose of naled (1NA Z=-3.03, p<0.01; PNPA Z=-6.05, p<0.01, Fig 5). Exposure to sublethal treatments of phenothrin or clothianidin did not significantly affect 1NA or PNPA activity 24 hours post treatment (Fig 5). Further application of lower sublethal doses of naled resulted in dose-dependent inhibition of 1NA (ρ=- 0.82, df=7, p=0.02) and PNPA activity (ρ=-0.75, df=7, p=0.05, Fig 6). Inhibition of PNPA activity was greater than inhibition of 1NA activity at 0.05 ug/bee (x^2^=12.5, df=1, p<0.01) and 0.066 ug/bee (x^2^=13.6, df=1, p<0.01; Fig 6).

**Fig 5.**
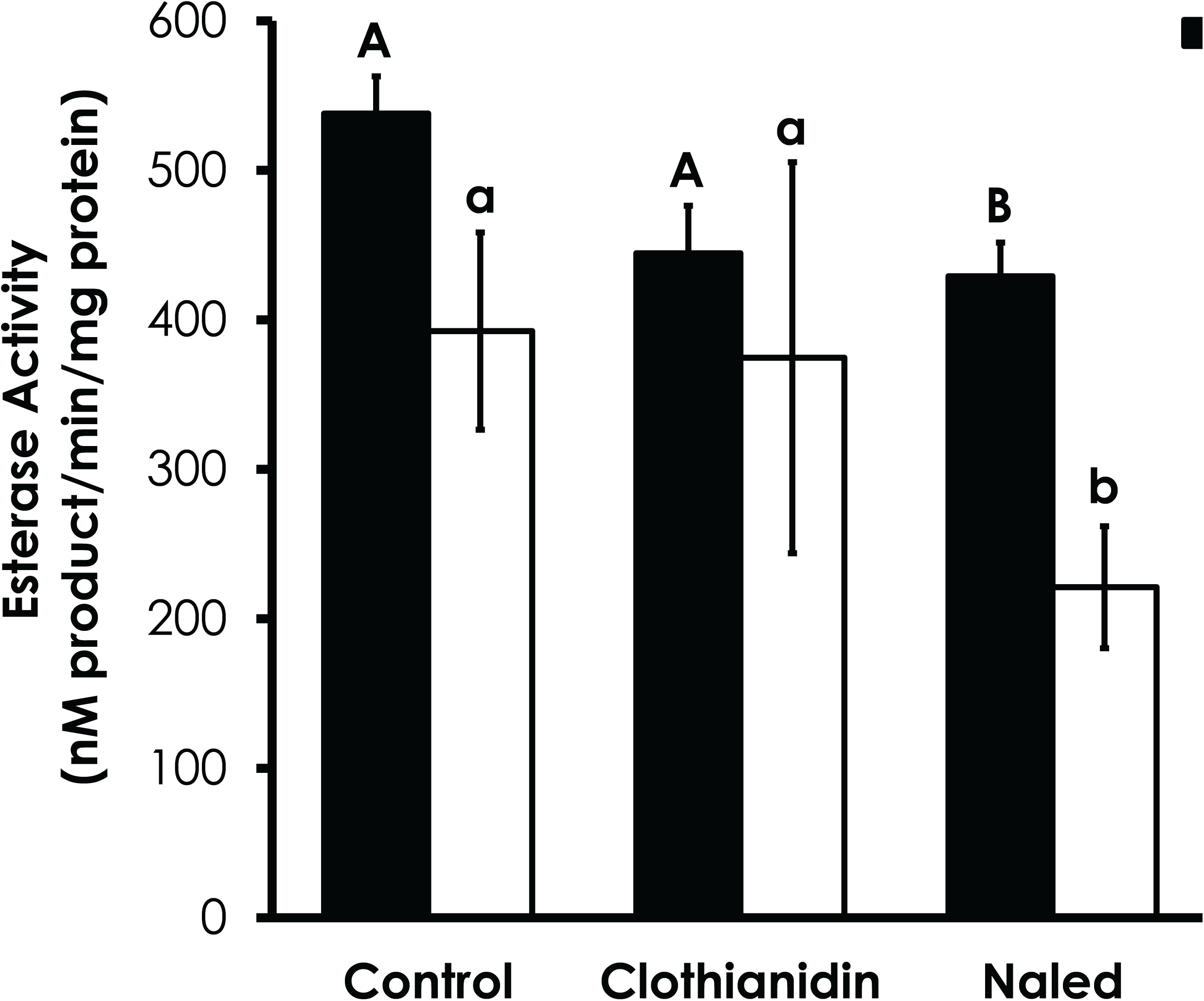
Effect of sublethal treatments of insecticides on esterase activity. Capital and lower case letters indicate significant differences in 1NA and PNPA activity, respectively

**Fig 6.**
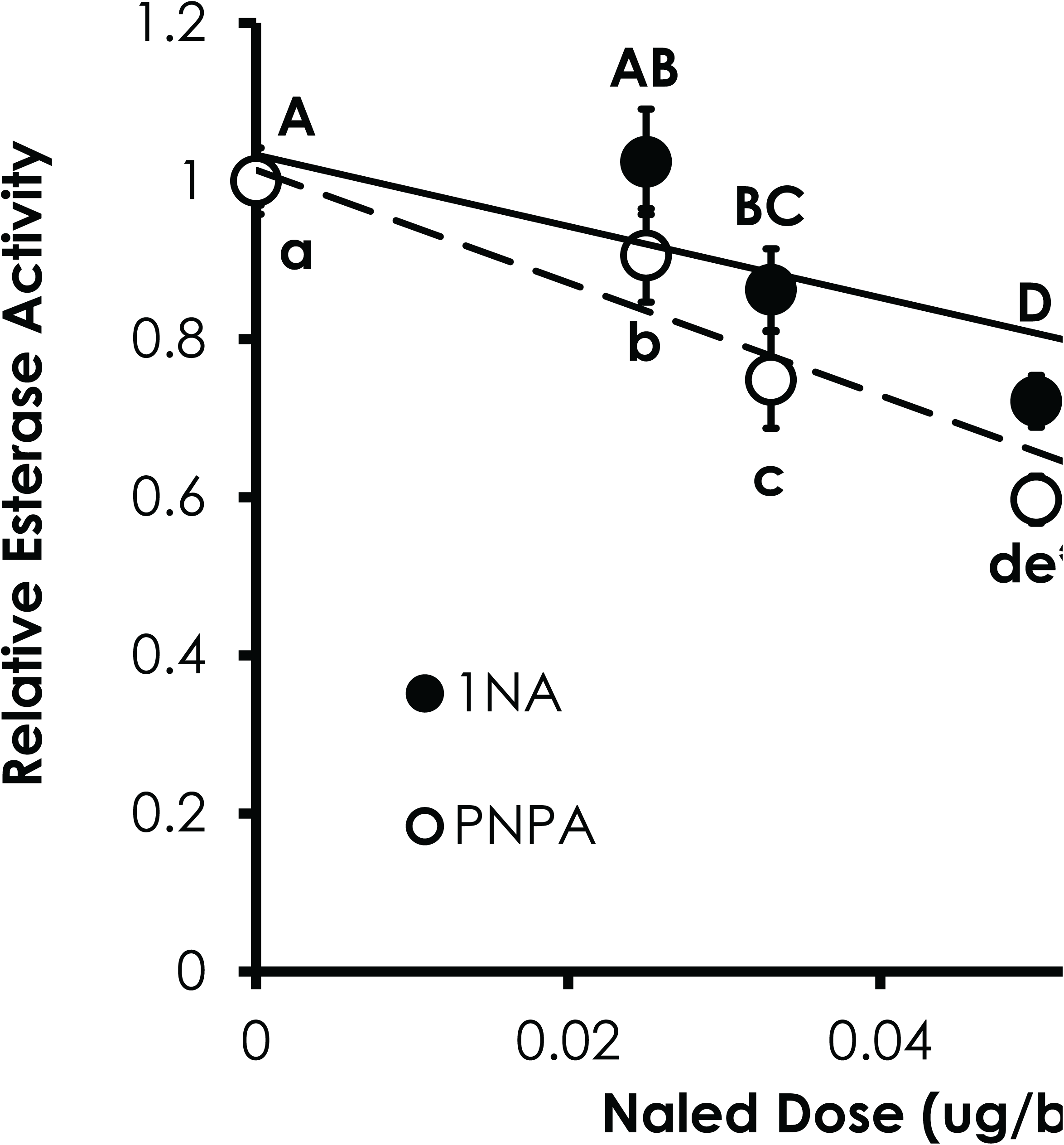
Relative inhibition of esterase activity towards 1NA and PNPA to sublethal doses of naled. Capital and lower case letters indicate significant differences in 1NA and PNPA activity, respectively. The asterisk indicate significant differences between substrates at respective naled doses.

### Viruses but not Varroa affect esterase activity

#### Varroa mite infestation does not affect esterase activity

Esterase activity towards 1NA (x^2^=0.30, df=1, p=0.58, Fig 7) or PNPA (x^2^=0.28, df=1, p=0.59, Fig 7) was not affected by Varroa mite infestation, as honey bee that pupated with a single Varroa mite feeding on them exhibited no differences in esterase activity compared to bees that developed without Varroa infestation.

**Fig 7.**
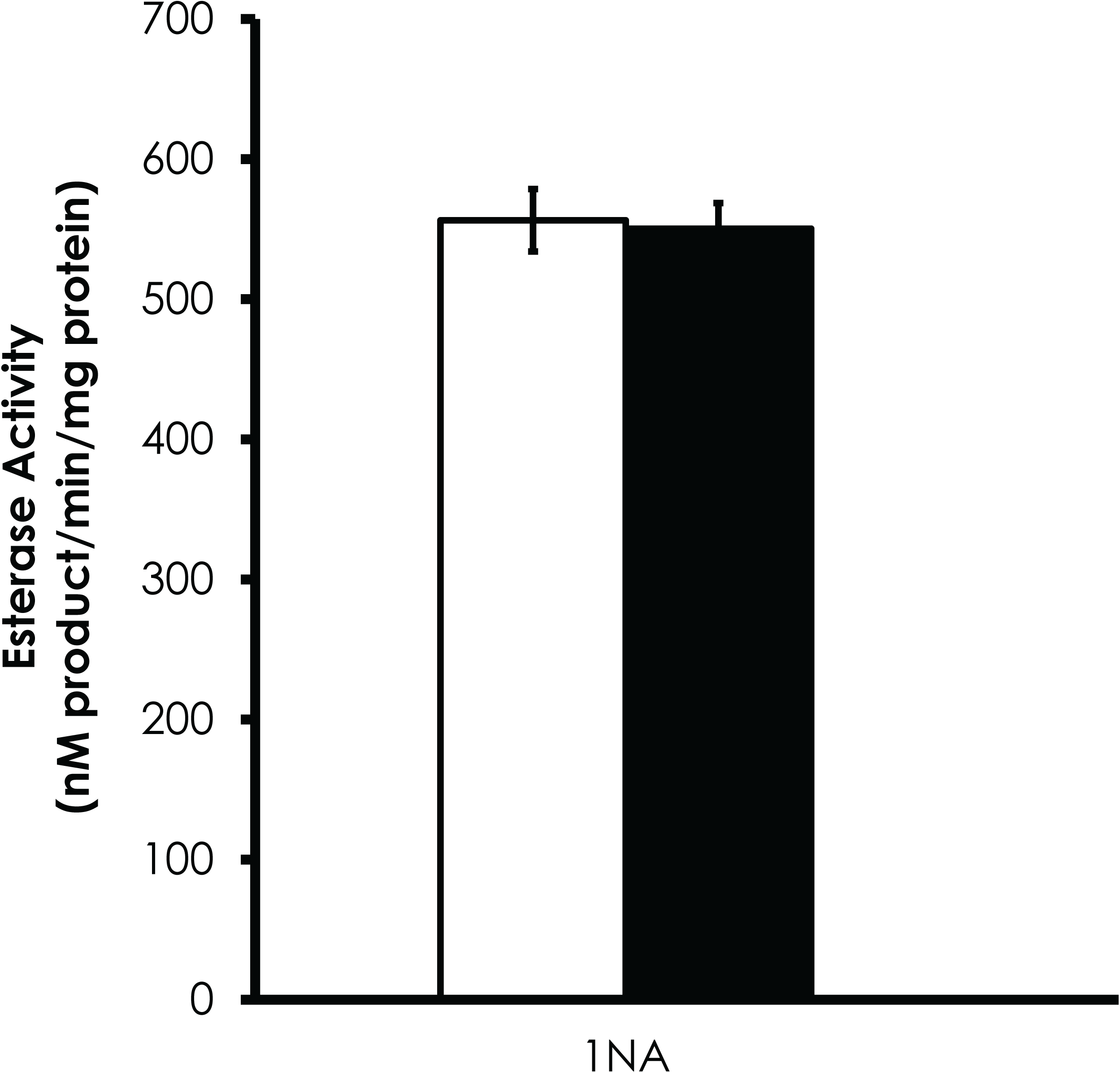
Varroa mite infestation does not affect esterase activity towards 1NA or PNPA. Data are shown as average ± SEM.

### Viral infection reduces esterase activity

Esterase activity towards 1NA significantly decreased in bees injected with BQCV, CBPV, and DWV relative to both uninjected and PBS-injected controls (F=18.8, df=12, p<0.01, Fig 8). PNPA activity was significantly reduced in bees injected with BQCV and CBPV compared to both uninjected and PBS-injected controls. DWV-injected bees had lower PNPA activity compared to uninjected controls (F=19.7, df=12, p<0.01, Fig 8).

**Fig 8.**
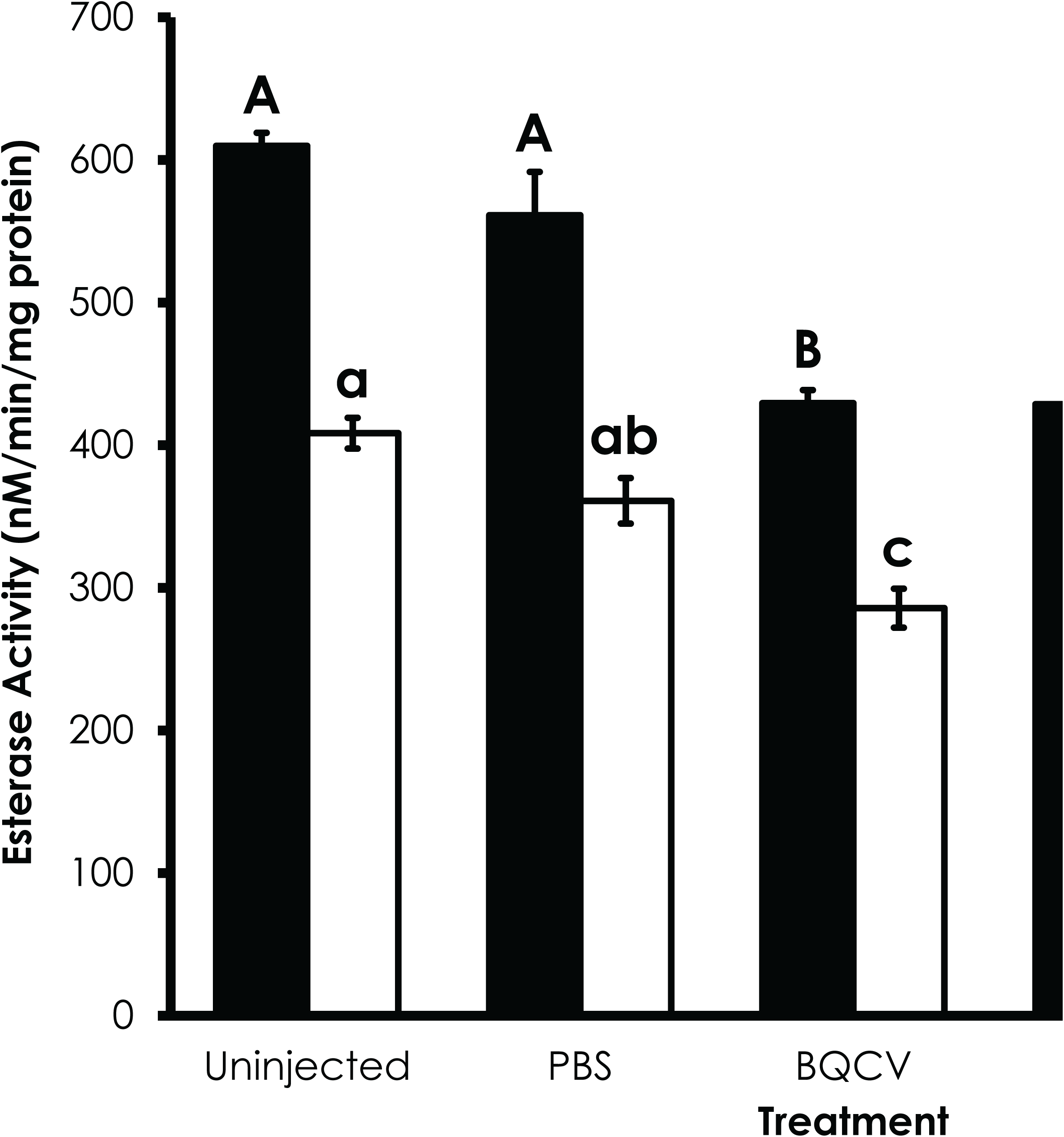
Viruses reduce esterase activity. Honey bees injected with BQCV (n=12), CVPV (n=16) or DWV (n=16) have significantly reduced esterase activity relative to uninjected (n=16) or PBS-injected controls (n=12). Capital and lower case letters indicate significant differences in esterase activity towards 1NA and PNPA, respectively. Data are the average ± SEM.

## Discussion

Our research shows plasticity in honey bee esterase activity due to a wide variety of life-history traits and external pressures. These physiological differences may underlie the wide range in the reports of honey bee pesticide sensitivity [16, 17] as well as emphasizing the need for detailed descriptions of the insecticide bioassay conditions in order to produce data that is comparable among researchers. The results we report on factors affecting esterase activity in honey bees are concepts that are easily applicable to other physiological systems such as immune function, nutrition utilization, and development where experimental conditions may dramatically affect the results. Honey bee colonies are complex and dynamic systems that are highly adaptable to changes in foraging resources, pathogen infection, parasite infestation, and pesticide exposure in particular. Understanding the physiological basis of how honey bees mediate these stresses allows for improved colony management strategies to promote honey bee colony health.

### Esterase activity does not correlate with insecticide sensitivity

We utilized 1NA and PNPA because they are model substrates that are indicative of general esterase and choline esterase activity, respectively [25, 40]. However, esterase activity towards the model substrates 1NA and PNPA may not reliable surrogates of esterase activity towards most insecticides in honey bee. It is possible that the activity of esterases capable of detoxifying organophosphates (OPs) cannot be assessed using the model substrates we employed. For example, a mutation in the E3 esterase of OP resistant strains of the sheep blowfly confers hydrolase activity towards the OP chlorfenvinphos while losing the ability to metabolize the model substrates 1NA and PNPA [8]. The difficulties of connecting insecticide sensitivity and esterase activity towards model substrates has been especially noted in OP resistant mosquitoes [41-43]. Thus, identification of honey bee-specific esterase substrates (including the insecticide itself) and inhibitors are urgently needed to accurately assess the metabolic contribution of esterases toward insecticide sensitivity.

The high esterase activity in Italian honey bees was an unexpected result since this stock of honey bees was the most sensitive to many insecticides and esterase inhibition produced the lowest level of synergism in phenothrin bioassays [13]. Comparison of esterase activity generated here with previously reported LD_50_/LC_50_ values among honey bee stocks [13] suggest variable roles for esterases in insecticide sensitivity. Although data points were limited in our previous study (and thus unable to be statistically analyzed appropriately), there was a positive association of esterase activity towards 1NA and PNPA with the LD_50_ of malathion among honey bee stocks. This is consistent with esterase-mediated detoxification of malathion [44]. There was no correlation with esterase activity towards 1NA and PNPA with the LD_50_ values of naled, etofenprox, resmethrin, or imidacloprid [13], suggesting other factors besides esterases are more important for explaining differential sensitivity to these insecticides among these honey bee stocks [45, 46]. Interestingly, esterase activity negatively correlated with the LD_50_s of phenothrin and thiamethoxam [13], suggesting that esterases may bioactivate these compounds to more toxic metabolites. However, the role of esterases in bioactivation of these compounds would be unusual as esterases are likely responsible for phenothrin detoxification [47], and P450s are responsible for bioactivation of thiamethoxam [48].

Comparison of the levels of esterase inhibition suggests a secondary role of esterases in determining phenothrin sensitivity. Italian honey bees had the highest levels of esterase activity towards PNPA but the lowest of level of synergism in phenothrin bioassays when the maximum sublethal dose of coumaphos was used to inhibit esterase activity [13]. However, this assumes that coumaphos provided similar levels of esterase inhibition among honey bee stocks and that coumaphos inhibits the esterases that are involved in phenothrin detoxification. Future studies on esterase inhibition with different inhibitors will help determine the types of esterase that contribute to activity towards these substrates and if there is any differences in the effectiveness of these inhibitors among honey bee stocks.

A third line of evidence that suggests a diminished role of esterases in insecticide detoxification is shown in the current study with the lack of correlation of esterase activity with phenothrin sensitivity and the negative correlation with naled sensitivity as bees aged. While esterases are important for phenothrin sensitivity in 3-day old bees [13, 14], the lack of correlation of esterase activity and phenothrin sensitivity with age suggests that other factors (i.e. P450s [14, 49]) may underlie the changes in phenothrin sensitivity with age. The negative association of esterase activity with naled sensitivity suggests esterase activity bioactivated naled. Bioactivation of OPs is typically accomplished via P450-mediated conversion of a thiophosphate to the active oxon species [50]. However, naled does not possess a thiophosphate. Therefore, bioactivation of naled by esterases is very unlikely due to its chemical structure. These results suggest that other factors besides esterases are important for determining the increased naled sensitivity with age in honey bees. Taken together, findings from current and previous work suggest that esterases activity as measured by metabolism of model substrates may play a secondary role in determining pesticide sensitivity [14].

### Esterase activity increases with honey bee age

Our results show that esterase activity increases with age in honey bees. This is consistent with the increase in cytochrome P450 and glutathione-S-transferase activities with age documented in honey bees [49, 51]. Sensitivity decreases to the pyrethroid, phenothrin, with age, while sensitivity increases to the OP, naled (Fig 3; [13]). Both P450s and esterases are involved in determining phenothrin sensitivity [13]. However, the lack of correlation of esterase activity with phenothrin sensitivity suggests that other factors, such as P450s, may be more important than esterases for causing the changes in phenothrin sensitivity with age. The increase in naled sensitivity with age is contradicted by the increase in esterase activity with age.

Previous work that showed that decrease in phenothrin sensitivity and an increase in naled sensitivity as honey bees aged in single cohort colonies, which are comprised of bees that are of the same chronological age but shift physiologically to conduct the different tasks needed for a functioning hive that would typically be divided across bee ages (e.g. feeding larvae vs. foraging) [13]. Those results are consistent with the results reported here for colonies with normal demographics (Fig 3). The similar changes in esterase activity and insecticide sensitivity with age in both types of colonies suggest that altered colony demographics do not affect insecticide sensitivity under our experimental conditions. It also suggests that chronological age is more significant than task (e.g. in-hive worker vs. forager) in determining sensitivity to insecticides, which is significant for toxicological bioassays.

### Esterase inhibition by insecticides

#### Clothianidin sensitivity

The LC_50_ value for clothianidin obtained with Italian honey bees was the same as the LC_50_ values for thiamethoxam reported in previous studies [13, 52]. Thiamethoxam must be bioactivated by cytochrome P450s [48] into clothianidin [53]. The similar LC_50_ values for thiamethoxam and clothianidin suggest that honey bees have a high metabolic capacity for this particular bioactivation. However, this phenomenon appears to be a common process as many other insects also possess similar LC_50_ values for thiamethoxam and clothianidin [53-57].

### Esterase inhibition is insecticide-dependent

The patterns of esterase inhibition by exposure to insecticides were expected based on their respective target sites. Naled is an OP that inhibits acetylcholinesterase activity, and most honey bee esterase activity is performed by choline esterases [40], therefore, it is not surprising naled inhibits esterase activity towards these model substrates. Since mortality occurs at doses of naled>0.066 ug/bee, it appears that inhibition of 26% and 42% of esterase activity towards 1NA and PNPA, respectively, results in mortality. Esterases significantly influence phenothrin sensitivity [13]. However, at the experimentally determined sublethal dose, phenothrin did not affect esterase activity towards 1NA or PNPA. Esterase activity was not affected by clothianidin exposure. This result is expected because studies on the effects of esterase inhibitors on sensitivity to clothianidin (or thiamethoxam) in honey bees have not been reported and these compounds have no ester bonds. Reports in which thiamethoxam (which is bioactivated *in vivo* to clothianidin [53]) has been shown to induce or inhibit esterase activity at concentrations near the LC_50_ [58], or at concentrations lower than the LC_50_ that would still result in low levels of mortality [11]. Our study indicates that esterases are not significantly inhibited *in vivo* at the much lower experimentally-determined sublethal concentrations of clothianidin that we employed here. While other studies have shown altered esterase activity to insecticide exposure at levels higher than the experimentally-determined sublethal levels we used here [11, 58], it is likely that mortality would be a more definitive and more convenient measure of insecticide exposure.

### Viruses transmitted by Varroa, but not Varroa infestation, impair esterase activity

Varroa infestation during the honey bees’ development did not alter esterase activity when they emerged as adults. Honey bees that have been infested by Varroa mites have smaller body size [32] and reduced expression of genes involved in metabolic detoxification [59, 60]. Our results, however, show no effect of Varroa mite infestation on esterase activity. This is consistent with the lack of change in insecticide sensitivity with varying Varroa mite infestation at the colony level [61]. Therefore, it appears that single-foundress Varroa infestation on its own may have little impact on pesticide sensitivity. However, the significant reduction in esterase activity by injection of virus shows that Varroa infestation can indirectly affect esterase activity as a disease vector. It is well known that viral infection can affect pesticide sensitivity [20, 24], and our results demonstrate that viruses may have significant effects on detoxification capacity. Future experiments with varying levels of viruses as well as focused investigation on the expression of genes involved in metabolic detoxification will demonstrate the impacts of viruses on honey bee health.

## Conclusions

This study demonstrates that honey bee esterase activity is very dynamic and significantly influenced by honey bee stock, age, insecticide exposure, and viral infection. Our results suggest a diminished or secondary role of esterases in determining insecticide sensitivity and that esterase activity toward model substrates does not accurately represent esterase activity towards insecticides. However, development of low-cost, high throughput assays using the insecticide as the esterase substrate would yield unambiguous results on the importance of esterases in determining insecticide sensitivity. The utility of using esterase activity towards model substrates as biomarkers of insecticide exposure should be pursued further and validated in order to be used as an accurate diagnostic tool. Despite these findings, reducing the quantity of insecticides as well as cautious and accurate application of insecticides near honey bee colonies as well as in foraging areas can reduce the potential negative impacts of insecticides on honey bee colony health. Besides potential effects from insecticides, honey bees are confronted with the significant and realistic problems of Varroa mites [62], introduced pathogens [63], loss of foraging area [64], and reduced queen health[65] and the complex interactions among all of these factors may work in concert to contribute to poor colony performance and productivity.

## Acknowledgements

We would like to thank Dave Dodge, Victor Rainey, Phil Tokarz, and Hunter Martin at USDA-ARS Honey Bee Breeding, Genetics and Physiology Laboratory for colony set up and maintenance, sample collection, virus purification and injection. This work was funded by a grant from the US Environmental Protection Agency (EPA Grant#83558201 to KBH) and the Louisiana State University Agricultural Center. This manuscript has been approved for publication by the Director of Louisiana Agricultural Experiment Station as Manuscript No 2016-234-28544. Mention of trade names or commercial products in this publication is solely for the purpose of providing specific information and does not imply recommendation or endorsement by the U.S. Department of Agriculture.

